# Permeabilization status affects the relationship between basal metabolic rate and mitochondrial respiration in great tit blood cells

**DOI:** 10.1101/2023.05.29.542691

**Authors:** Elisa Thoral, Carmen C. García Díaz, Elin Persson, Imen Chamkha, Eskil Elmér, Suvi Ruuskanen, Andreas Nord

## Abstract

Although mitochondrial respiration is believed to explain a substantial part of the variation in whole-animal basal (BMR) or resting metabolic rate (RMR), few studies have addressed the relationship between organismal and cellular metabolism and how this may vary in environments where individual demands for energy differ. We investigated the relationship between whole-individual metabolic rate, measured in temperatures ranging thermoneutrality to far below thermoneutrality, and mitochondrial respiration of intact or permeabilized blood cells in two separate studies on wild great tits (*Parus major* L.). Our results show that, in permeabilized cells, there are significant positive relationships between BMR or RMR and several mitochondrial traits, including phosphorylating respiration rate through both complexes I and II (i.e., OXPHOS respiration). However, surprisingly, the LEAK respiration (i.e., basal respiration that mainly counteract for proton leakage) was not related to BMR or RMR. When measurements were performed using intact blood cells, BMR was positively related to ROUTINE respiration (i.e., mitochondrial respiration on endogenous substrates) in one of the two studies, but no other mitochondrial traits could explain variation in BMR or RMR in any thermal environment. These studies seem to show that the level of activation of mitochondrial metabolism as well as the permeabilization status of blood cells play a primary role on the extent to which blood metabolism might explain variations in the whole-individual metabolic rate.

## Introduction

Energy spent by individuals to fuel physiological and behavioural processes can be estimated by measuring metabolic rate, *i.e.* the oxygen consumed per unit time (Norin and Metcalfe, 2019). In endotherms, the minimum energy consumption needed to sustain life at rest in a post-absorptive state and in thermoneutrality is defined as the basal metabolic rate (BMR), or resting metabolic rate (RMR) if the measurement is performed outside of the thermoneutral zone (Bligh and Johnson, 2001). This RMR varies across environmental, temporal, and spatial contexts. Accordingly, tropical birds are known to have a lower RMR compared to temperate species (Wiersma et al., 2007), while migrant avian species that mainly live in cold habitats at high latitude locations have a higher RMR (Jetz et al., 2008). On smaller geographic scales, RMR varies predictably with environmental conditions such as temperature (McKechnie and Swanson, 2010; White et al., 2007). Hence, many birds that winter in the temperate zone show seasonal upregulation of RMR to meet the higher demands for energy processing in the wintertime (Swanson and Vézina, 2015; Dawson and Yacoe, 1983). Thus, by influencing the idle cost of living, RMR or BMR potentially set the limits for several fundamental evolutionary processes, including pace of life, reconciliation of the trade-off between energy acquisition and loss (Londoño et al., 2015; Wikelski et al., 2003), and the amount of energy amenable for investment into performance (aerobic scope, Buttemer et al., 2019; Dutenhoffer and Swanson, 1996), all of which can be linked to fitness (but see Arnold et al., 2021). This, in turn, can explain why RMR or BMR typically has high heritability (e.g. Nilsson et al., 2009; but see McFarlane et al., 2021) and is subjected to differential selection depending on environmental context (Nilsson and Nilsson, 2016). Thus, understanding the proximal drivers of variation in metabolic rate in natural populations is essential to better understand how environmental conditions shape metabolic phenotypes.

At the cellular level, the vast majority of energy that is subsequently used to fuel RMR and BMR is produced by oxidative phosphorylation (OXPHOS) in mitochondria (Lehninger et al., 1993). Mitochondria use oxygen and energy substrates to transport electrons from complex I to IV (cytochrome c oxidase) of the electron transport system (ETS) whilst simultaneously forming a proton gradient across the inner mitochondrial membrane. Chemical energy in the form of adenosine triphosphate (ATP) is formed when this proton motive force is dissipated through complex V (ATP synthase) at the end of the ETS (Lanza and Nair, 2009). However, the coupling between oxygen consumption and ATP production is not complete, such that a part of mitochondrial respiration is used to counteract proton leakage back into the mitochondrial matrix (i.e., LEAK respiration). This causes energy contained in the proton gradient to be released as heat and is the basis for non-shivering thermogenesis in endothermic animals, studied mostly in brown adipose tissue of mammals (e.g. Jastroch et al., 2010). Moreover, the ETS also leads to the production of reactive oxygen species which are responsible for oxidative stress if they are present in too large quantities in the body (Munro and Treberg, 2017). Thus, the coupling efficiency of the ETS, i.e., the balance between substrate oxidation and phosphorylation, vary with environmental conditions met by individuals, notably thermal environment. For example, an increase in LEAK respiration is observed in adult zebra finches (*Taeniopygia guttata*) that experienced early-life heat waves (Ton et al., 2021), possibly as a potential protective mechanism against oxidative stress. Moreover, after several weeks at 5°C, an increase in LEAK respiration is also observed in liver of tree shrews (*Tupaia belangeri*), potentially to improve non-shivering thermogenesis in the cold (Zhang et al., 2011). In the same way, LEAK respiration in red blood cells of several small birds species is higher in colder winter compared to mild autumn temperature, which also reduces the coupling of oxygen consumption and phosphorylation but likely increases mitochondrial thermogenesis (Nord et al., 2021). Because metabolic rate at the whole-animal and cellular levels seem to respond similarly to environmental perturbations, mitochondrial metabolism could be a good candidate pathway to explain the mechanistic underpinnings of variation in organismal metabolism (Koch et al., 2021; Swanson et al., 2014; White and Kearney, 2013). In line with this, RMR is positively related to the LEAK respiration and the activity of cytochrome c oxidase in several tissues (including muscle, liver and kidney) in Chinese hwameis (*Garrulax canorus*) (Wang et al., 2019). In black-capped chickadees (*Poecile atricapillus*), BMR is positively correlated with phosphorylating respiration rate (i.e., OXPHOS respiration) in the liver of cold-acclimated birds (Milbergue et al., 2022). Moreover, in great tits (*Parus major*), RMR is positively related to endogenous mitochondrial metabolism in intact blood cells, provided birds are not stressed (i.e., have lower circulating levels of corticosterone), but weakly negatively related to mitochondrial respiration when birds are stressed (Malkoc et al., 2021).

Mitochondrial respiration and ETS function are rarely static. For example, while LEAK respiration may explain some 20-25% of BMR in mammals under constant conditions (Porter and Brand, 1995; Rolfe and Brand, 1996), other studies show that there is significant variation in LEAK respiration in relation to, for example, metabolic and thermogenic demand and in response to environmental stressors (Divakaruni and Brand, 2011; Roussel and Voituron, 2020; Speakman et al., 2004; Talbot et al., 2004). Thus, it can be speculated that the slope, sign, and intercept of the relationship between organismal and cellular respiration can change depending on both individual state and environmental context. We tested this hypothesis using the “little bird in winter” model – i.e., a small (< 25 g) endothermic and homeothermic songbird with high surface area to volume ratio and high metabolic intensity that face considerable thermogenic costs all the while enduring difficult foraging prospects due to short days, cold temperature, and low winter food availability (Brodin, 2007). Specifically, we investigated the relationship between whole-animal BMR and RMR, measured in different night-time temperatures ranging thermoneutrality to far below thermoneutrality, with mitochondrial metabolism of blood cells in winter-acclimatised wild great tits (*Parus major* L.). The red blood cells of birds and most other non-mammalian vertebrates contain functional mitochondria (Moras et al., 2017; Moritz et al., 1997), allowing insight into mitochondrial metabolism and function without the need for killing the animals to collect tissues, which is often necessary for mitochondrial studies in other tissues. Blood cell measurement in birds is also feasible on minute volumes that can be safely extracted from small organisms under field conditions (Nord et al., 2021; Stier et al., 2019). As blood is, therefore, a very promising tissue to study, we were interested in exploring to which extent blood cell mitochondria can predict whole-animal physiology. For this purpose, we measured metabolic rate overnight in a range of different environmental temperature-conditions and then performed mitochondrial measurements the morning after, using both intact cells respiring only on endogenous (intracellular) substrates and permeabilized cells in the presence of saturating concentrations of mitochondrial substrates and adenosine diphosphate (ADP). This allowed us to study the link between organismal and cellular metabolism both in representative *in vivo* conditions (intact cells) and at the level of ETS function when not constrained by substrate availability (permeabilized cells). Our purposes were two-fold: (i) to investigate the extent to which blood cell respiration is indicative of organismal metabolic rate, for which there is currently mixed evidence, and (ii) to test experimentally if the contribution of blood mitochondrial respiration and respiratory complexes towards explaining variation in organismal BMR and RMR changes depending on environmental context of the individuals.

## Material and Methods

Data used in the study originate from two separate experiments on wild great tits where organismal metabolic rate and mitochondrial respiration in blood cells were measured as part of different sub-projects. Upon capture, age and sex were determined based on plumage characteristics, and body mass (± 0.1g) was recorded before and after the measurement of organismal metabolic rate.

### Study 1 – Intact blood cells

A detailed description of bird capture, instrumentation and experimental protocol can be found in García-Díaz et al., 2023. Briefly, great tits (n = 57) were captured in mist nets at sunflower-baited bird feeders during daytime, or when roosting at nest boxes during night-time, between January and February 2021 at two sites in southernmost Sweden (Räften: 55° 43’N, 13° 17’E”); Vomb: 55° 39’N, 13° 33’E). The birds were brought to the animal housing facilities at Lund University within 2 h of capture and were instrumented intraperitoneally with a temperature-sensitive PIT tag under local anaesthesia for use in a different study. The birds were housed singly in cages measuring 77 × 44 × 77 cm (width × depth × height) under simulated natural photoperiod at 5°C, with *ad libitum* access to food (sunflower seeds, peanuts, lard) and water. The birds were released at their capture site once metabolic rate measurements had been completed.

### Whole-individual metabolic rate

We measured metabolic rate using flow-through respirometry during night-time between 12 and 36 h after capture. Birds were alternately assigned to 3 groups for metabolic rate measurement in different thermal conditions: (i) a simulated cold winter night (mean temperature ± SD: −15.12 ± 0.15°C), (ii) a simulated normal winter night (5.17 ± 0.03°C) or (iii) in thermoneutrality (24.93 ± 0.10°C). The birds were put into a 1L hermetically sealed glass container ventilated with dry (drierite) air at 400-500 mL/min (standard temperature and pressure, dry, STPD; registered using a FB-8 mass flow meter from Sable Systems, Las Vegas, NV, USA) and contained inside a climate chamber (Weiss Umwelttechnik C180 [Reiskirchen, Germany]). This flow rate was sufficient to keep CO_2_ in outgoing air below 0.5% (measured using a CA-10 carbon dioxide analyser; Sable Systems). Temperature inside the respirometry chambers was recorded using a TC-2000 thermocouple box (Sable Systems International, Las Vegas, USA) fitted with fine-wire (36 G) type T (copper-constantan) thermocouples with the temperature-sensitive junction contained in the chamber roof where they were minimally affected by heat production by the bird. The fraction of oxygen in outgoing air was measured in dry, CO_2_-free (drierite, ascarite) air using a FC-10 oxygen analyser (Sable Systems) and was converted to metabolic rate assuming an energy equivalence of 20 J per mL O_2_ (Kleiber, 1961). Each bird was measured in one temperature only.

### Mitochondrial measurements

We collected a blood sample for mitochondrial measurements from the jugular vein (≤ 10% of total blood volume) within 1 min of removing the birds from the metabolic chambers in the morning after RMR/BMR measurements. The sample was contained in 2mL K_2_-EDTA (dipotassium ethylenediaminetetraacetic acid) tubes (BD Vacutainer^®^, Franklin Lakes, NJ, USA) that were kept at 10-12°C until further processing. Thirty minutes to 2 h later, we measured the mitochondrial metabolism in intact blood cells based on whole blood measurement at 41°C (a representative daytime body temperature for a passerine bird; Prinzinger et al., 1991) using high-resolution respirometers (Oxygraph O2k high-resolution respirometers, Oroboros Instruments, Innsbruck, Austria) following Nord et al., 2023.

After mixing, 50µL whole blood was added to 1.95 mL of respiration medium (MiRO5: 0.5mM EGTA, 3mM MgCl_2_, 60mM K-lactobionate, 20mM taurine, 10mM KH_2_PO_4_, 20mM HEPES, 110mM sucrose, 1g/L free-fatty-acid bovine serum albumin, pH 7.1) in the respirometer chambers. We first measured baseline respiration rate, which is directly related to the oxygen consumption on endogenous substrates (*i.e.,* ROUTINE respiration). Oligomycin (chamber concentration: 5µM) was then added to inhibit ATP synthase. Since ADP cannot be phosphorylated to ATP in the presence of oligomycin, the remaining oxygen consumption is mostly used to counteract for proton leakage across the inner mitochondrial membrane (*i.e.,* LEAK respiration). After that, the maximal respiration rate of the ETS (*i.e.,* ETS respiration) was stimulated by titration of the mitochondrial uncoupler cyanide-p-trifluoro-methoxyphenyl-hydrazone (FCCP). The ETS respiration was defined as the maximum oxygen consumption measured after a first injection of 1.5μL of FCCP (solution concentration: 1mM) followed by subsequent injections of 0.5µL until maximum. Finally, antimycin A (5µM) was added to inhibit complex III, after which any remaining respiration is of non-mitochondrial original (*i.e.,* residual oxygen consumption, or ROX respiration). ROX was removed from all other respiration rates before analyses.

### Study 2 – Permeabilized blood cells

Great tits (n = 73) were captured from December 2021 to March 2022 when roosting in nest boxes at night in three sites within close proximity in southernmost Sweden: Linnebjer (55° 43’N 13° 17’E; n = 7), Frueräften (55° 72ʹN, 13° 31ʹE; n = 8) and Vomb (55° 39’N, 13° 33’E; n = 59). Linnebjer and Frueräften are adjacent to the Räften woodlot sampled in Experiment 1 and all have similar habitat characteristics. The Vomb site was the same as in Experiment 1. The birds were returned to the nest box in which they had been caught at, or just after, sunrise the next day.

### Whole-individual metabolic rate

The birds were brought to Lund University within 3 h of capture and their BMR was measured firstly in thermoneutrality (20.33 ± 0.84°C) for 4 h and then the RMR was measured below thermoneutrality at 8°C (7.98 ± 0.18°C) for 4 h using the respirometry and climate chamber setup described above. The temperature decrease was 6.25°C per h during the transition between the warm and mild temperature. We sub-sampled 150 mL/min STPD from the main gas stream (which was 400-500 mL/min STPD) using a SS4 sub-sampler (Sable Systems). O_2_ was measured in dry, but not CO_2_-free air, and so we mathematically accounted for the effect of CO_2_ on O_2_ when calculating metabolic rate (Lighton, 2019).

### Mitochondrial measurement

The birds were bled (≤ 10% of blood volume) from the jugular vein within 1 min of removal from the metabolic chambers in the morning after BMR/RMR measurements, and the sample was stored in 4 mL Na-heparin tubes (BD Vacutainer^®^, Franklin Lakes, NJ, USA) at 10-12°C until processed 30 minutes to 2.5 h later. Samples from 12 birds caught in November 2021 were measured as in Study 1 (*i.e.*, in intact blood cells). The blood metabolism of the remaining 61 individuals was recorded in permeabilized blood cells. All measurements were done at 41°C. Briefly, 40µL of blood were added to 1mL of ice-cold MiR05 and mixed gently. Then, 50µL from this mixture was transferred to a new tube containing 950µL of PBS and stored at 5°C for subsequent the cell counting (using a TC20 Automated Cell Counter, Bio-Rad, Solna, Sweden) 10 to 30 min later (when the mitochondrial measurements had started). Cells were counted in an attempt to normalize the mitochondrial respiration. However, since there were no relationships between mitochondrial respiration traits and cell count when two clearly outlying observations were excluded from the analyses (see Figure S1), these data were not considered further.

The blood-MiR05 tube was centrifuged for 2 min at 1000 RCF and the supernatant was removed. The pellet was then resuspended in 1mL of 41°C MiR05 from the respirometry chamber and the entire solution was put back in the chamber to reach a final volume of 2.1 mL. Once ROUTINE had been measured, digitonin (40µg/mL) was added to permeabilize the cells. Once oxygen consumption had stabilised after digitonin addition, we added malate (chamber concentration: 5mM) and pyruvate (2mM) to fuel complex I of the ETS to obtain a basal oxygen consumption (*i.e.,* Background respiration). Then, we stimulated the production of ATP through complex I (*i.e.,* OXPHOS_CI_) by adding ADP (1.25mM). When the response had stabilised, we stimulated complex II by adding 10 mM succinate to obtain maximal phosphorylating respiration rate (*i.e.,* OXPHOS_CI+II_). LEAK respiration was measured by injection of oligomycin (2.5µM), after which ROX was obtained by addition of antimycin A (2.5µM). As in Experiment 1, ROX respiration was removed from all the other respiration before analyses.

### Data handling and statistical analyses

In Study 1, to study the relationship between whole-individual metabolic rate and functional traits of the mitochondria, we calculated two flux control ratios (FCRs): E-R control efficiency (FCR_R/E_), an index of the maximum working capacity of the ETS used during an endogenous respiration; and E-L coupling efficiency (FCR_L/E_), an index of the mitochondrial efficiency to produce ATP in a stimulated cellular state (Gnaiger, 2020).

In Study 2, also to relate metabolic rate to functional aspects of mitochondrial respiration, we calculated the Net Phosphorylation Efficiency, which is an index of the mitochondrial efficiency to produce ATP by oxidative phosphorylation (Shama et al., 2016). The calculation method and definition of the different respiration rates and FCRs are detailed in Table 1.

**Table 1:**
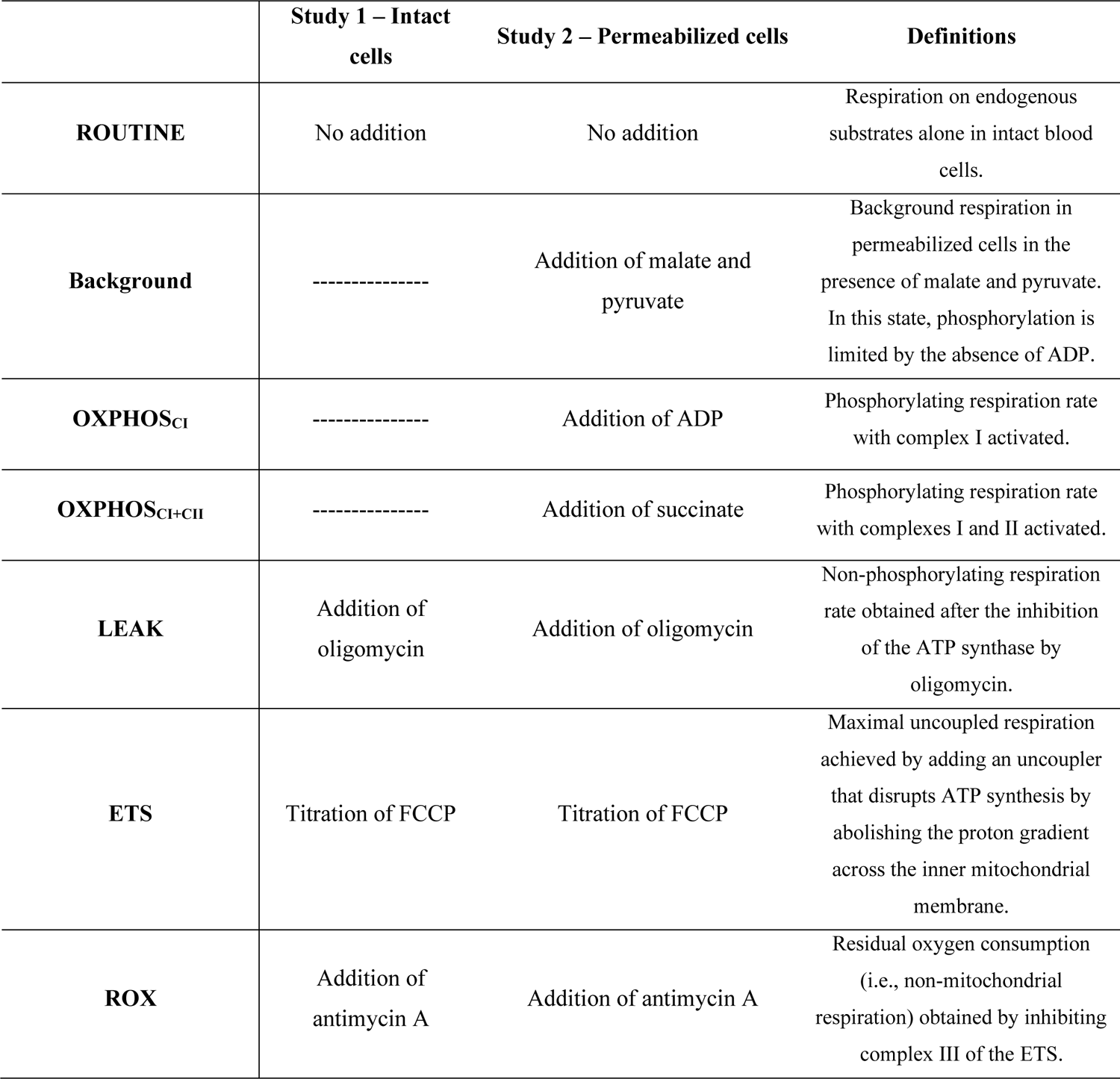
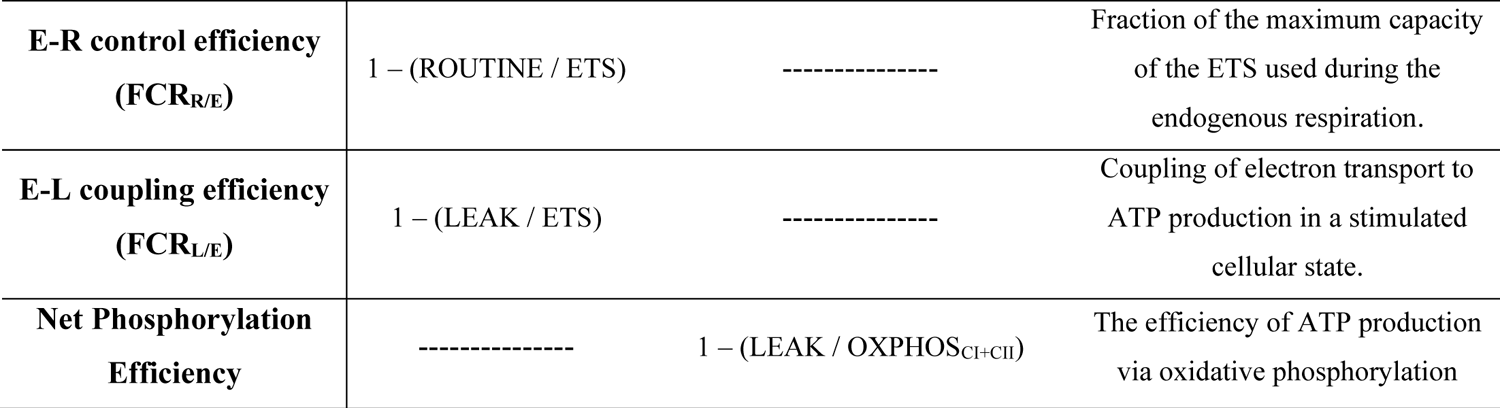
Description of mitochondrial respiration parameters, flux control ratios (FCRs) and their derivation when relation cellular metabolism to organismal basal or resting metabolic rate. Data were collected in two separate studies of wild, winter-acclimated great tits. In Study 1, mitochondrial measurements were performed using intact blood cells in whole blood samples, whereas in Study 2, the blood cells were permeabilized and measured in the presence of saturating amounts of mitochondrial substrates.

We combined the data obtained on intact cells and metabolic rate measurement at 20°C in Study 2 with those from the thermoneutral group (n = 12, *i.e.,* +25°C) in Study 1 since both are well within the thermoneutral zone of great tits (lower critical temperature: 12.9°C; AN, unpublished data). In Study 2, we only present the relationships between BMR and mitochondrial parameters in the main document, because the slopes of the relationships between metabolic rate and mitochondrial respiration traits never differed significantly (respiration trait × temperature treatment: all *P* ≥ 0.2). Accordingly, final models showed at most minor differences (Table 3 and Table S1), and data for RMR measured at 8°C are presented in the Supplementary Data (Figure S2).

We excluded all data from one bird with an infected wound in Study 1, to avoid any effects of an ongoing immune response on organismal or cellular respiration (Ots et al., 2001; Van De Crommenacker et al., 2010). At the mitochondrial level, we also excluded samples where either oligomycin or ADP did not affect mitochondrial respiration, when overall respiration was close to 0 or when the signal was not stable at all during the measurement (n = 9 of 57 in Study 1 [4 from cold winter night condition, 3 for normal winter night condition, 2 from thermoneutrality]; n = 2 of 11 for the intact cells and n = 10 of 61 for the permeabilized cells in Study 2). We also removed two BMR measurements in Study 2 since the gas consumption curves clearly indicated that the birds did not conform to the “resting criterion” of BMR (Bligh and Johnson, 2001; BMR > population mean + 2 standard deviations). Data on Background respiration in Study 2 are missing for 6 individuals since ADP was added directly after pyruvate and malate, preventing the measurement of Background respiration alone.

All statistical tests were done using R *v.* 4.2.1 (R Development Core Team 2022). In both studies, linear models (lm in R base) were fitted to estimate the effects of mitochondrial respiration parameters (ROUTINE, LEAK and ETS respirations for Study 1; ROUTINE, Background, OXPHOS and LEAK respirations and Net Phosphorylation Efficiency for Study 2) on BMR and RMR. We included body mass as a covariate in all models since it is often an important predictor of metabolic rate. Body mass and mitochondrial parameters were mean-centered (within temperature-treatment groups) in both studies, using the *isolate()* function from the *bmlm* package (Vuorre and Bolger, 2018). In both experiments, we fitted separate models for each temperature at which the metabolic rate was measured, and for each mitochondrial parameter. The latter was necessary since the variance inflation factor (VIF) was in the range of 2-7 when all temperatures were analysed in single models. All data met the assumptions of normality and homogeneity of variance as determined by examination of diagnostic plots of the final models. The level of significance was set at *P* ≤ 0.05. All analyses were performed with and without outliers (i.e., observations that deviated ≥ 2 standard deviations from the group mean). Relations with outliers are presented in Supplemental Data (Study 1: Figure S3, Study 2: Figure S4).

## Results

All parameters of the linear regression models are presented in Table 2 (Study 1) and Table 3 (Study 2). In both studies, RMR is used to describe the metabolic rate measured below thermoneutrality whereas BMR is used to describe the metabolic rate obtained at thermoneutrality.

**Table 2:**
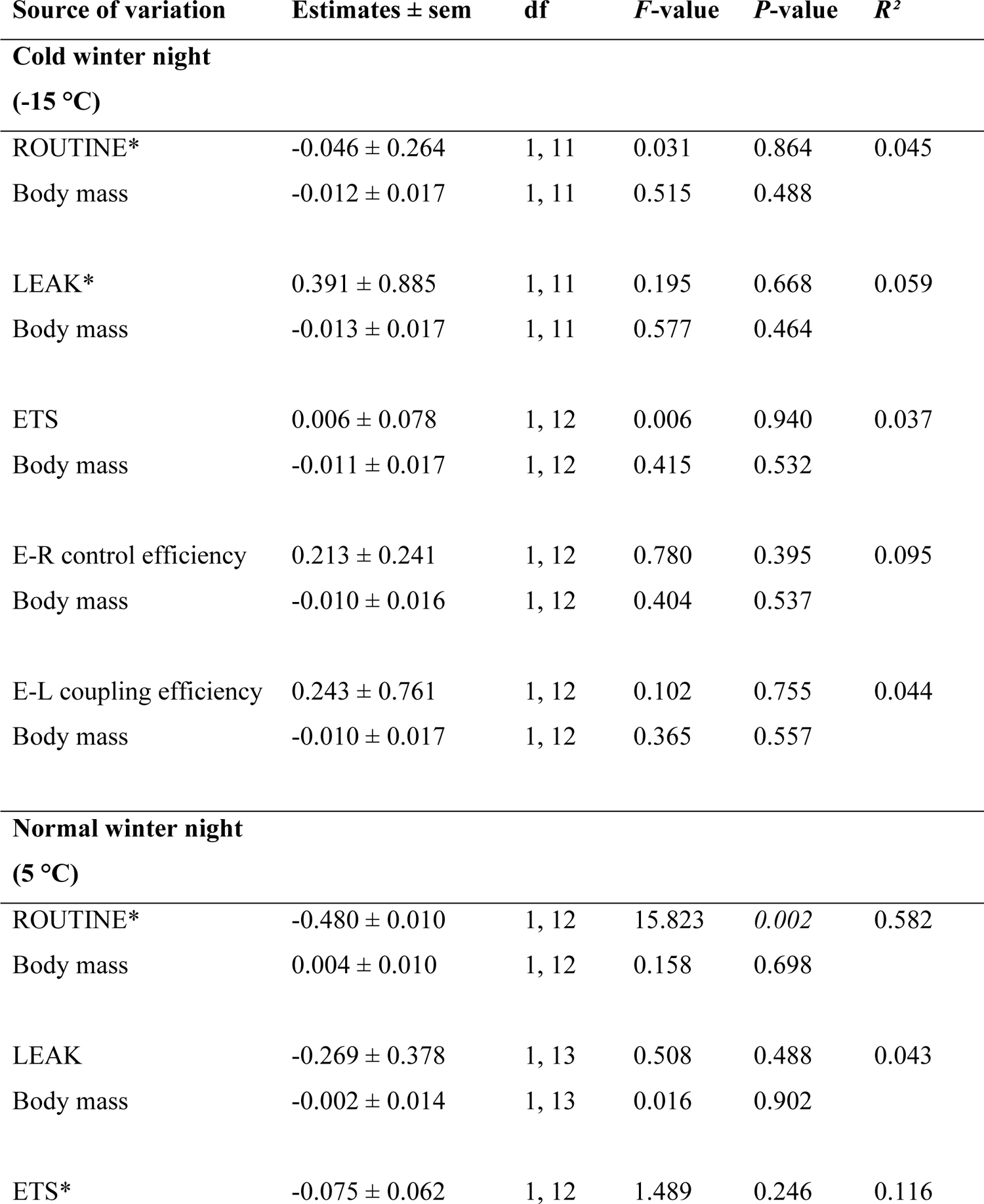
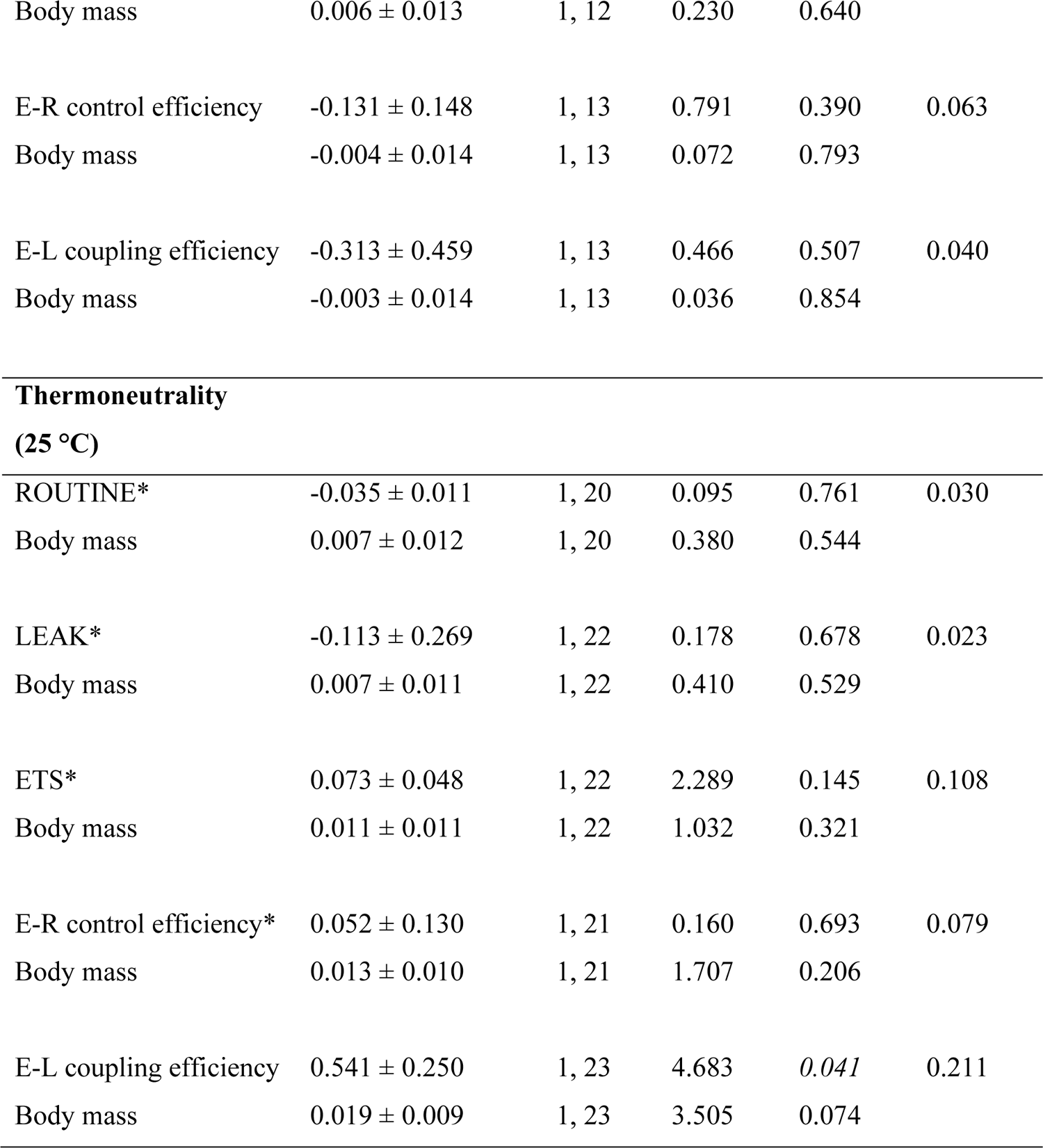
Parameter estimates from linear models of the relationship between resting or basal metabolic rate (RMR or BMR) and mitochondrial metabolism in intact blood cells of wild great tits in Study 1 (and Study 2 at thermoneutrality). Body mass and mitochondrial parameters were group-mean centered before analyses. Significant *P*-values (i.e., *P* ≤ 0.05) are presented in italic font. *Parameters for which one or several outliers have been removed.

**Table 3:**
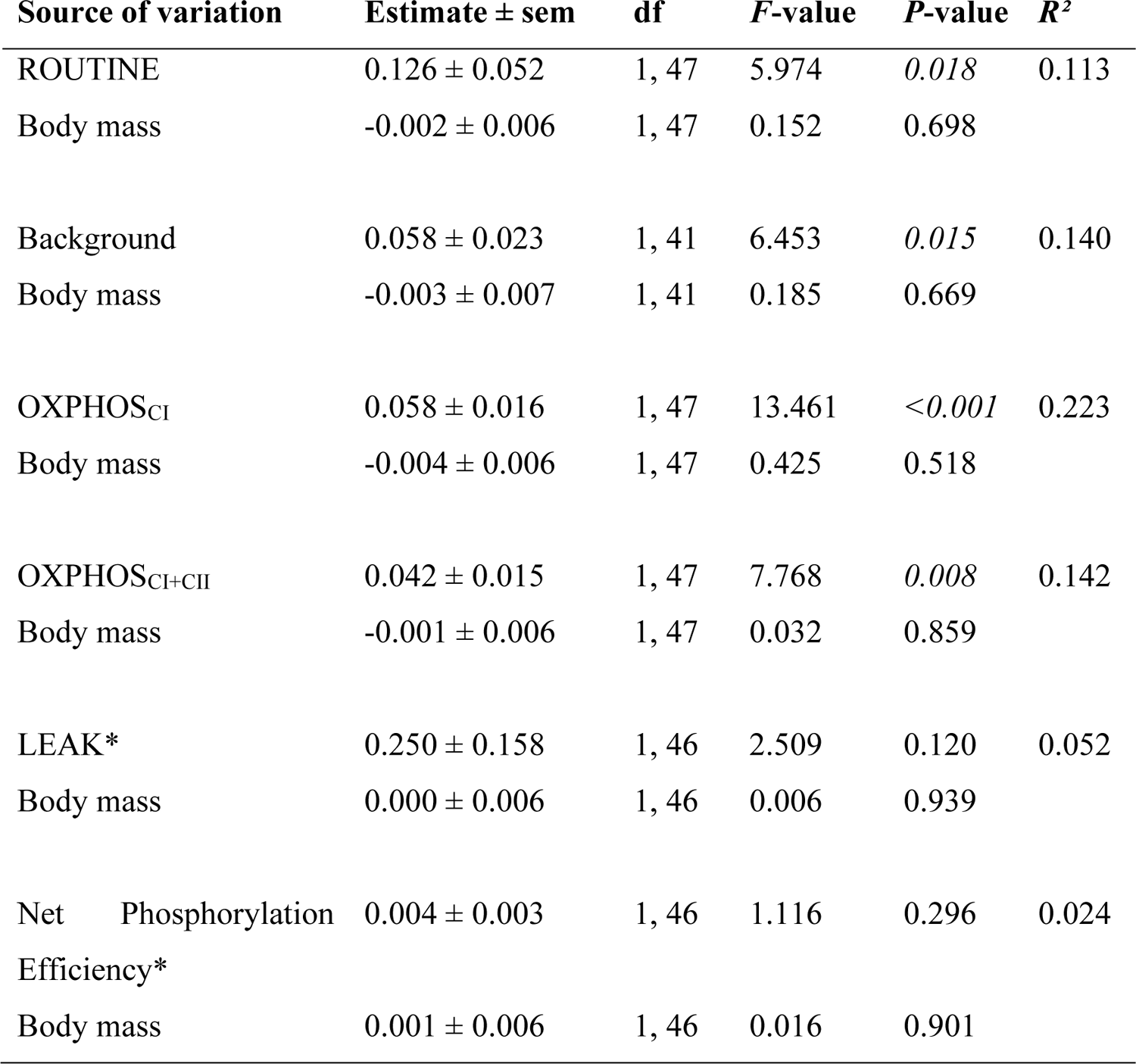
Parameter estimates from linear models of the relationship between BMR and mitochondrial metabolism in permeabilized blood cells of wild great tits in Study 2. Body mass and mitochondrial parameters were group-mean centered before analyses. *P*-values in italics are significant (i.e., *P* ≤ 0.05). *Parameters for which one outlier has been removed.

### Relationship between BMR or RMR and mitochondrial respiration in intact blood cells

There was a significant negative relationship between RMR measured in normal winter night conditions (*i.e.,* in 5°C) and ROUTINE respiration, whereby RMR declined by 48W for each unit increase in ROUTINE (*P* = 0.002, Figure 1A, Table 2), but we found no such relationship in cold winter night conditions (*P* = 0.864) or in thermoneutrality (*P* = 0.761). However, there was a significant positive relationship between BMR and E-L coupling efficiency at thermoneutrality (*i.e.,* FCR_L/E_; *P* = 0.041, Figure 1E, Table 2), that was not found during cold (*P* = 0.755) and normal winter night conditions (*P* = 0.507). No other respiration traits or flux control ratios explained any significant part of the variation in BMR or RMR (all *P* > 0.1; Figure 1; Table 2).

**Figure 1:**
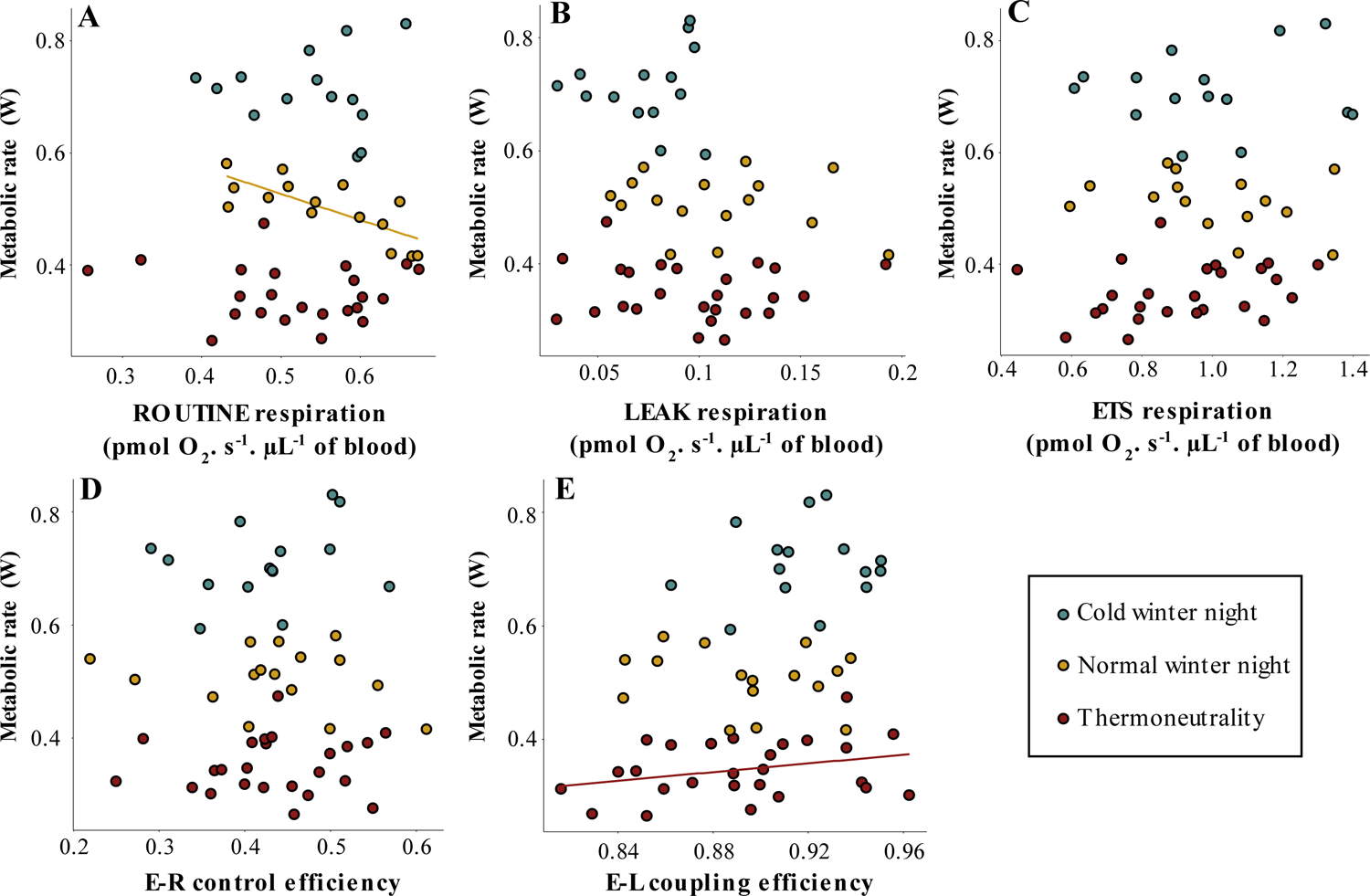
Relationship between RMR or BMR and mitochondrial parameters in intact blood cells. Data were collected from wild, winter-acclimated great tits. Mitochondrial measurements were performed on intact cells in whole blood samples, where the mitochondria were respiring only on endogenous substrates in different night temperatures. A single bird was measured in one night temperature condition only. The solid line indicates a significant relationship (*P* ≤ 0.05), and absence of a regression line indicates that the relationship was non-significant.

### Relationship between BMR or RMR and mitochondrial respiration in permeabilized blood cells

In Study 2, when measurements were performed using permeabilized cells, we found that variation in BMR was significantly explained by several mitochondrial parameters. In contrast to Study 1, ROUTINE respiration before permeabilization was positively related to BMR (*P* = 0.018, Figure 2A, Table 3), such that BMR increased by 13W for each unit increase in ROUTINE. Moreover, there were significant positive relationships between BMR and Background respiration (*P* = 0.015, Figure 2B, Table 3) and OXPHOS respiration through both complex I and complex I+II (OXPHOS_CI_: *P* < 0.001, OXPHOS_CI+CII_: *P* = 0.008, Figures 2C and 2D, Table 3), whereby BMR increased by 5.8, 5.8 and 4.2W for each unit increase in Background, OXPHOS_CI_ and OXPHOS_CI+CII_ respirations, respectively. However, neither LEAK respiration and Net Phosphorylation Efficiency were related to BMR (LEAK: *P* = 0.120, Figure 2E, Table 3; Net Phosphorylation Efficiency: *P* = 0.296, Figure 2F; Table 3).

**Figure 2:**
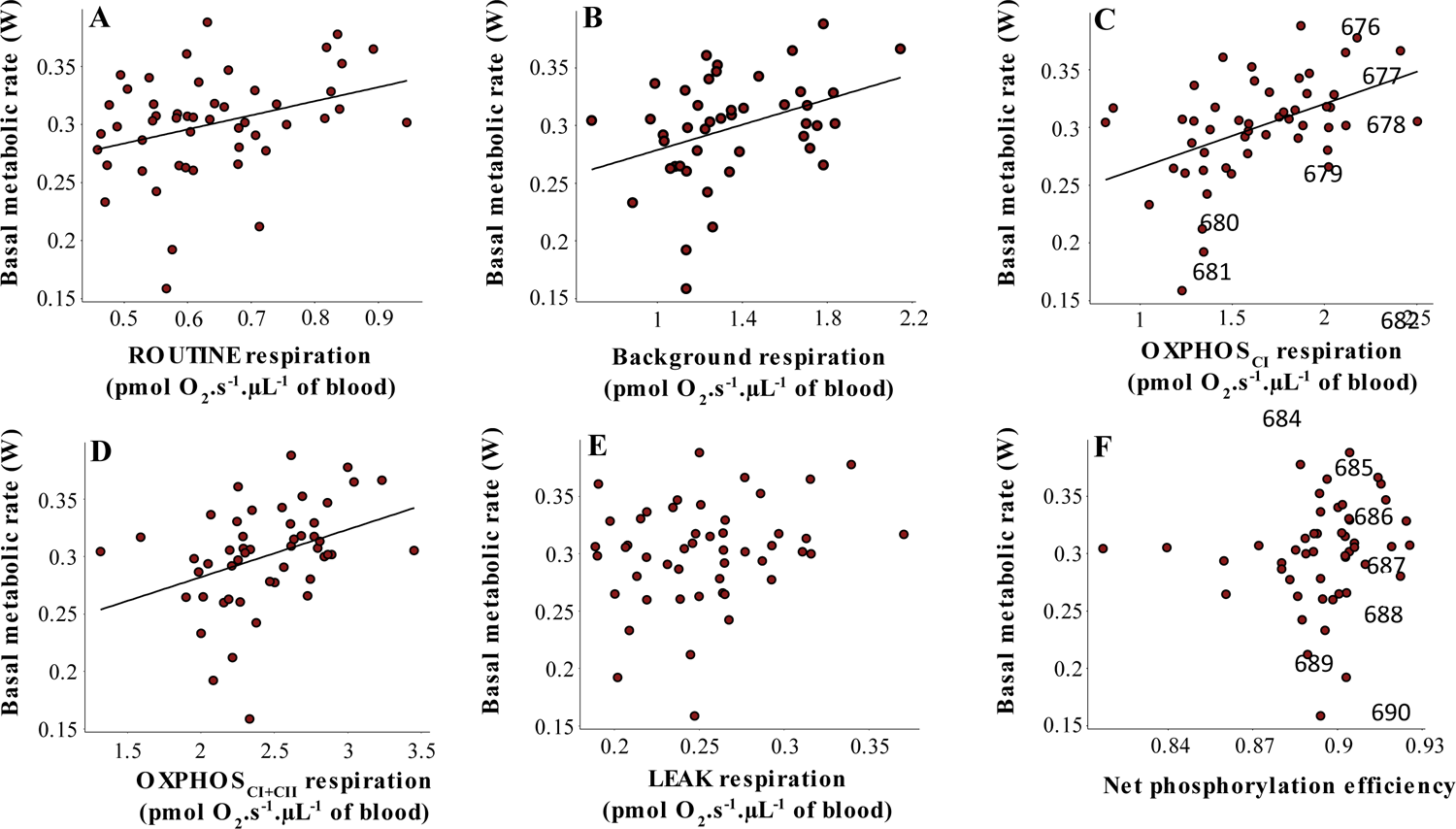
Relationship between BMR and mitochondrial respiration parameters in permeabilized blood cells. Data were collected from wild, winter-acclimated, great tits. Apart from ROUTINE (A), all respirations were obtained after permeabilization and after addition of different substrates or inhibitors of the ETS (see Table 1). Solid lines show significant relationships (*P* ≤ 0.05) whereas absence of a regression line indicates non-significant relationships.

The relationships between mitochondrial parameters and RMR measured at 7.5°C were similar to those for BMR, except for the Background respiration which was no longer significant (Figure S2 Panel 1B, Table S1).

### The effects of outliers

Figures with outliers are presented in Figure S3 for Study 1 and in Figures S2 Panel 2 and Figure S4 for Study 2. In Study 1, keeping outliers in the model did not change the overall results, except in the case of ROUTINE respiration which was no longer significantly related to RMR at 5°C (*P* = 0.146, Figure S3A, Table S2) and the ETS respiration that tended to be related to RMR at 5°C after including one outlier (*P* = 0.053, Figure S3C, Table S2). In Study 2, keeping outliers did not change any relationships at thermoneutrality (Figure S4, Table S3) or below thermoneutrality (Figure S2, Table S1).

## Discussion

The results obtained in our two studies indicate that the permeabilization level of red blood cells as well as the activation level of the mitochondrial metabolism seem to play a major role in the relationship between the blood metabolism and whole-individual metabolic rate. In contrast, the environmental temperature experienced when measuring the whole-individual metabolic rate globally did not affect this relationship.

### Permeabilization status of red blood cells affects the relationship between mitochondrial metabolism and individual metabolic rate

Apart from a positive effect of the E-L coupling efficiency (FCR_L/E_; Figure 1E) and a negative effect of the ROUTINE respiration that will be detailed later (Figure 1A), individual metabolic rate was not significantly explained by any mitochondrial respiration traits measured in intact blood cells. However, when the cells were permeabilized, BMR was significantly positively related to several mitochondrial parameters. A possible explanation for the effect of permeabilization status could be the provision of unlimited amounts of substrates in permeabilized cells bypasses any rate-limiting steps that may influence mitochondrial function (Djafarzadeh and Jakob, 2017). For example, LEAK respiration in intact cells in our study was only 37% and 8% of the LEAK and Background respirations, respectively, in permeabilized cells. Similarly, maximal respiration in intact cells (*i.e.,* ETS) was only 44% of that in permeabilized cells (*i.e.,* OXPHOS_CI+II_). Thus, it can be speculated that we cannot directly measure changes in mitochondrial physiology in response to environmental change in intact cells, where respiration rate is limited by substrate availability (Kyriazis et al., 2022; Leverve, 2007), transmembrane transport rate (Osellame et al., 2012), and any interactions between mitochondria and other cell components (Boldogh and Pon, 2007; Wieckowski et al., 2009). However, it must be noted that environmental conditions under which metabolism was studied differ at the cellular and whole-individual levels. Indeed, while the conditions under which mitochondrial metabolism was measured in intact blood cells approximated those experienced by the birds at the time of whole-individual metabolic rate measurement, i.e., no unlimited access to food or energy substrates, only significant correlations were measured between whole-individual metabolic rate and mitochondrial metabolism measured in permeabilized cells, i.e., in with saturated amounts of energetic substrates. Thus, although blood cell permeabilization could be a necessary step to directly study the link between mitochondrial metabolism and whole-individual metabolic rate, one should keep in mind that the conditions for measuring mitochondrial metabolism cannot be achieved at the individual level.

### Activation level of the electron transport system affects the relationship between mitochondrial respiration and individual metabolic rate

In permeabilized cells, several metrics of OXPHOS respiration were positively related to metabolic rate (Figures 2C and D; Figure S2C and D). Thus, in blood cells, the respiration directly associated with energy production seems to contribute to whole-individual metabolic rate. Similar results were observed by Milbergue et al., (2022) where OXPHOS_CI_ measured in liver cells was positively correlated to RMR at −10°C. Several studies suggest a role of blood cell mitochondria in oxidative stress and ageing (Delhaye et al., 2016; Jimenez et al., 2019), which could be physiologically linked to the intensity of ATP production (Chung and Schulte, 2020; Koch et al., 2021). Another hypothesis is that ATP produced in the blood could participate in the maintenance of basal functions of vascularized tissues, so that the level of OXPHOS respiration is proportional to the amount of metabolizing tissues. However, no study has formally addressed the role of ATP produced by blood cells, making this an exciting avenue for further investigation in the future.

Surprisingly, LEAK respiration was never related to BMR, which contrasts studies in mammals where LEAK has been found to explain some 30% of the variation in BMR (Brand et al., 1999; Rolfe et al., 1999). On the other hand, studies on humans found no relationship between LEAK and BMR and instead suggest that metabolic rate is determined more by the mitochondrial oxygen affinity and density (Larsen et al., 2011). Moreover, the relationship between mitochondrial and organismal respirations may also vary among tissues. For example, seasonal variations in the size of muscle and excretory organs, such as liver and kidneys, are responsible for changes in BMR in black-capped chickadees (*Poecile atricapillus*) (Petit et al., 2014). Moreover, in the same species, BMR was correlated to LEAK respiration in skeletal muscle, but not in the liver (Milbergue et al. 2022). On the contrary, standard metabolic rate in brown trout (*Salmo trutta*) was correlated to LEAK in the liver, but not in the muscle (Salin et al., 2016). Thus, in contrast to the phosphorylating capacity, LEAK respiration in the blood was not related to whole-individual metabolic rate in great tits. Indeed, the capacity to dissipate energy in blood might be more associated with an enhanced whole-individual metabolic rate, such as maximum metabolic rate, as observed in white muscle from brown trout (Salin et al., 2016). However, unlike LEAK respiration, Background respiration was significantly and positively related to BMR (Fig. 2B). ATP synthase is not inhibited when measuring Background respiration and the mitochondria are fuelled only by complex I substrates. In the LEAK state, ATP synthase is inhibited by oligomycin and both complexes I and II are activated. While it seems obvious that any intracellular ADP will be much diluted in the respiration medium after permeabilization, the only difference between these two respirations is then the activation or not of complex II. Then, it might be possible that, at a basal activation level of the ETS, the activity of complex I might be more related to the individual metabolic rate compared to complex II. To test this hypothesis, it would be interesting to measure the basal activity of complex II by blocking complex-I activity, thanks to an inhibitor such as rotenone, and assess the relationship between complex-II activity and the whole-individual metabolic rate.

### What does blood cell metabolism tell us about whole-individual metabolic rate?

On a more general note, we do not know if the observed significant relationships between metabolic rate and OXPHOS respiration in the permeabilized blood cells were causal or correlated. The latter could be the more likely since blood cells are not very aerobic compared to, for example, liver and muscle. Perhaps mitochondrial respiration in blood cells is representative of an overall mitochondrial phenotype that is correlated with organismal metabolic rate depending on the physiological and environmental conditions in which the relationship is studied? It should be kept in mind, however, that the temperature at which the RMR or BMR was measured had, under our conditions, little effect on the relationship between individual metabolic rate and blood metabolism. One possibility would be that too rapid changes in temperature (few hours) could not lead to changes in mitochondrial metabolism in blood, either by intrinsic changes in mitochondrial functioning, or by the increased presence of a new blood cell phenotype better acclimated to the new environment (Voss et al., 2010). Thus, future studies measuring the effect of a longer exposure to different temperatures on the relationship between individual metabolic rate and blood metabolism would directly measure the role of environmental temperature in modulating metabolism at different biological levels. It is also interesting to note that the relationship between metabolic rate and ROUTINE respiration differed between studies. Accordingly, during a simulated normal winter night in Study 1 (*i.e.,* at +5°C), ROUTINE respiration was *negatively* related to RMR (Figure 1A), as observed in Malkoc et al., (2021) for birds with high circulating levels of glucocorticoids. However, the ROUTINE respiration measured before permeabilization in Study 2 was *positively* related to BMR (Figure 2A), similarly to what was reported for low-glucocorticoid phenotypes in Malkoc et al. (2021). This could be related to the number of individuals per group in Study 1 that was smaller than in Study 2, leading to more inter-individual variations. It is also possible that the inversion of this relationship could be explained by interannual differences in mean physiological trait levels, such as in the amount of hormone secretion, that could impact metabolic rate via non-mitochondrial pathways (Williams, 2008; Williams, 2012). Alternatively, and non-exclusively, these differences might also be explained by variations in experimental conditions experienced by the birds. Specifically, in Study 1, birds were caught during daytime and kept in captivity for up to 36 h before being measured. Therefore, these birds were exposed to new environmental conditions and other interventions prior to sampling which could increase stress levels. In Study 2, birds were caught in the wild when roosting in nest boxes at night and their metabolic rate was measured within hours of capture. Thus, they had been eating natural food and had been subjected only to natural conditions before the experiment started, perhaps making them more similar to the “low-stress” phenotype in the study by Malkoc et al., (2021). Future studies should also assess the physiological mechanisms that mediate this switch of sign in the relationship between organismal and cellular metabolism.

## Acknowledgements

The authors thank Camilla Björklöv for assistance with animal care in Study 1, and Stefan Nord for managing the Räften feeders where birds for Study 1 were caught.

## Author contributions

AN and ET conceived the ideas and designed the methodology together with EE, IC, and SR; CCGD, EP, IC and AN collected the data. ET analysed the data; ET and AN led the writing of the manuscript. All authors contributed critically to the drafts and gave final approval for publication.

## Data availability statement

The data will be made available once the article is accepted for publication.

